# Variation in behavior drives multiscale responses to habitat conditions in timber rattlesnakes (Crotalus horridus)

**DOI:** 10.1101/2020.12.03.410290

**Authors:** Andrew S. Hoffman, Annalee M. Tutterow, Meaghan R. Gade, Bryce T. Adams, William E. Peterman

## Abstract

Variations in both the behavior of wildlife and the scale at which the environment most influences the space use of wild animals (i.e., scale of effect) are critical, but often overlooked in habitat selection modeling. Ecologists have proposed that biological responses happening over longer time frames are influenced by environmental variables at larger spatial scales, but this has rarely been empirically tested. Here, we hypothesized that long-term patterns of behavior (i.e. lasting multiple weeks to months) would be associated with larger scales of effect than more sporadic behaviors. We predicted site use by 43 radio-telemetered timber rattlesnakes (*Crotalus horridus*) exhibiting four distinct, time-varying behaviors (foraging, digestion, ecdysis, and gestation) using remotely-sensed environmental variables related to forest structure and landscape topography. Among sites used by snakes, warmer temperatures and higher levels of forest disturbance were predictive of behaviors dependent on thermoregulation including gestation and ecdysis while more moderate temperatures and drier, more oak-dominated sites were predictive of foraging. Long-term behaviors were associated with larger spatial scales, supporting our hypothesis that the scale at which habitat selection occurs is linked to the temporal scale of relevant behaviors. Management recommendations based on single-scale models of habitat use that do not account for fine-scale variations in behavior may obscure the importance of potentially limiting habitat features needed for infrequent behaviors that are critically important for growth and reproduction of this and related species.

## Introduction

Organisms respond to environmental variation simultaneously across numerous temporal and spatial scales. Understanding habitat use across these scales is key to developing appropriate management strategies for rare or threatened wildlife. Modern telemetry and GIS technology allow for the collection of increasingly high-resolution data and can pair well with traditional point selection function (PSF) analyses (Boyce et al. 2002, Zeller et al. 2014). More recently, an increasing number of studies have stressed the importance of scale in such analyses, and have highlighted that few studies empirically select appropriate scales for habitat models (Holland et al. 2004, Smith et al. 2011, Jackson and Fahrig 2015, McGarigal et al. 2016). Empirical testing of appropriate scales, especially the grain of environmental variables, is critical given the recent proliferation of and ready access to remotely-sensed data that can help explain wildlife site use (Thompson and McGarigal 2002, Neumann et al. 2015).

The appropriate spatial extent at which to measure habitat features is the “scale of effect” (Jackson and Fahrig 2012) and depends on species’ traits and the biological variable of interest (Miguet et al., 2016). Miguet et al. (2016) discuss a framework for assessing the scale of effect for different variables that might influence a species’ space use. The authors assert that biological processes occurring over longer time periods (e.g., occurrence) will have larger spatial scales of effect than those happening over short time periods (e.g., fecundity). Although relatively few studies have found support for this hypothesis (Cushman and McGarigal 2004, Jackson and Fahrig 2014), the general idea of a link between temporal and spatial scales is a foundational principal of landscape ecology (Peterson et al. 1998).

A temporal-spatial scale connection should be evident within the temporally variable behavioral patterns of wildlife. For ectotherms, site-specific behavior is often related to thermoregulation, which drives much of their habitat selection (Stevenson 1985, Huey 1991). Both the thermal landscape and energetic requirements of ectotherms fluctuate seasonally, contributing to variation in habitat use throughout the year (Waldron et al. 2006, George et al. 2017). However, individuals within a single population often operate under different selective pressures at the same time of year, owing to variation in body condition or reproductive state (Lesmerises and St-Laurent 2017). Thus, generalizing population-level, seasonal shifts in behavior and site use may overlook important, fine-scale heterogeneity.

Behavioral variation is often difficult to assess when direct observation is infrequent or may disrupt natural behaviors. Rattlesnakes are ideal subjects with which to study the effect behavior has on habitat selection as they can be directly observed using radio-telemetry with minimal disturbance (Brown et al. 1982, Reinert and Zappalorti 1988) and they exhibit stereotyped behaviors (Clark 2004, Reinert et al. 2011a). While previous studies have detailed the effect that gestation has on space use in snakes (Charland and Gregory 1995, Reinert and Zappalorti 1988, Sprague and Bateman 2018), other behaviors have been largely ignored. Waldron et al. (2006) identified “seasons” based on foraging, breeding, and hibernation to describe space use in timber rattlesnakes (*Crotalus horridus*) but did not address changes in behavior within a season.

Rattlesnakes are ovoviviparous with long (4–5 months), energetically-costly gestation periods (Ernst and Ernst 2003). Females reduce movements, seek out relatively warmer or more exposed sites than would normally be used, and decrease or stop foraging for food (Brown 1991, Martin 1993, Gardner-Santana and Beaupre 2009). Ecdysis and digestion occur over a shorter timeframe (7–14 days and 3–7 days respectively) but these physiological states can also drive snakes to select more open habitats for thermoregulation (Greenwald and Kanter 1979, Semlitsch 1979). Foraging behavior may be even more sporadic and opportunistic, with associated changes in behavior during a single day (Clark 2004, Reinert et al. 2011a). Snakes making opportunistic site use decisions related to foraging and digestion may consequently select habitat at finer scales than snakes making decisions related to more predictable, long-term behavioral states (i.e. gestation and ecdysis).

We tested whether predictable, long-term behavior patterns have a larger scale of effect when compared to more sporadic, short-term behaviors in timber rattlesnakes. We considered a multi-scale framework to determine the appropriate scale of effect of several geospatial landscape features on four behaviors of interest in wild timber rattlesnakes. We hypothesized that snakes making short-term decisions (i.e. foraging or digestion) on site use would select habitat at finer scales than snakes making decisions related to predictable, long-term behaviors (i.e. gestation or ecdysis).

## Methods

### Study Site

Vinton Furnace State Experimental Forest (VFSF) is a 4,892-ha property located in Vinton County, Ohio, USA, and within the Southern Unglaciated Allegheny Plateau Ecological section (Cleland et al. 2007). The landscape features a dissected topography, including sharp ridges and valleys with relatively low relief (~100 m). The surrounding region is primarily forested, consisting of mostly second-growth stands recovering from heavy exploitation following the mid-to-late 1800s (Stout, 1933). The mixed-mesophytic forest type of the region is primarily dominated by oak species (*Quercus* spp.), especially on dry ridgetops and southwestern hillslopes, and transitions to mesophytic forest assemblages, including sugar maple (*Acer saccharum*), American beech (*Fagus grandifolia*), and yellow poplar (*Liriodendron tulipifera*), on opposing northeastern hillslopes and bottomlands (Hix and Pearcy; Adams et al. 2019). The study site also includes small tracts of pine plantations, usually composed of monospecific stands of *Pinus strobus*, *P. resinosa*, or *P. echinata*, while other native conifers, including *P. virginiana*, *P. rigida*, and *Tsuga canadensis*, combine at low densities with deciduous hardwoods. In addition, the study site incorporates active, ongoing forest management, especially within the central Raccoon Ecological Management Area. Varying management approaches, including even- and uneven-aged silvicultural prescriptions, as well as prescribed fire, promote heterogeneity in habitat conditions within the study site (Ducey 1982).

### Telemetry

We captured individual timber rattlesnakes and surgically implanted intraperitoneal transmitters (Holohil SI-2T) following procedures outlined by Reinert and Cundall (1982), using machine-administered isoflurane to anesthetize animals via inhalation. Transmitter weight did not exceed 5% of the snake’s body weight, and in most instances was < 2%. We released snakes at their original capture location within 24 hours of surgery and relocated individuals 2–3 times per week from April through October 2016–2019. Upon visually locating a snake, we recorded the location with a Global Positioning System (Garmin GPSmap 64s) to an estimated < 5m spatial accuracy.

### Behavior Classification

We identified four behavioral and physiological states (hereafter referred to as “behaviors”): foraging, digestion, ecdysis, and gestation. We considered snakes to be foraging when they were observed in a stereotyped foraging posture (Reinert et al. 2011a). Our observations of snakes digesting a recent meal are limited by the difficulty in detecting a small bolus in a large-bodied snake, but we noted this whenever possible. These observations are, therefore, inherently biased toward snakes digesting larger, more obvious meals and may be prone to a high false negative detection rate. We noted observations of snakes in ecdysis as indicated by snakes having blue-gray eyes and dusky skin coloration. We confirmed observations of gestating females by sonogram shortly after snake emergence in April and May. We excluded all locations in which a snake was not visible and presumably underground or actively moving. When not moving and not clearly participating in the aforementioned behaviors, we designated snake behavior as resting. We excluded snake observations from analyses when there was uncertainty about behavioral classification.

### Geospatial Habitat Features

We incorporated 15 geospatial habitat features, characterizing vegetation structure, plant species composition, and environmental variation, at varying resolutions (5-30 m), in our analysis of multiscale rattlesnake habitat use (Table 1). Digital canopy height (CHM) and elevation (DEM) models were developed from LiDAR data provided by the Ohio Geospatial Reference Program, collected in 2007 (http://ogrip.oit.ohio.gov/; accessed 13 October 2014). The LiDAR data featured two returns pulse^−1^ with an average spacing and density of 1.27 m and 0.27 returns m^−2^, respectively. The CHM and DEM models incorporated conventional methods, using bilinear interpolation, at 5 m resolution. From the DEM, we developed layers on slope (°), solar radiation, and the Beer’s transformed aspect (an index ranging 0–2, corresponding to southwestern to northeastern aspects, respectively). In addition to canopy height represented within the CHM, we developed three penetration ratios (number of returns <2 m in height divided by the number of returns <50 m and <10 m, including returns <2m; and the number of returns <1 m divided by the number of returns <5 m, including returns <1 m) to characterize overstory, midstory, and understory vegetation density at 30 m resolution (Adams and Matthews, 2019; Müller et al. 2010, Melin et al. 2018).

**Table 1.**
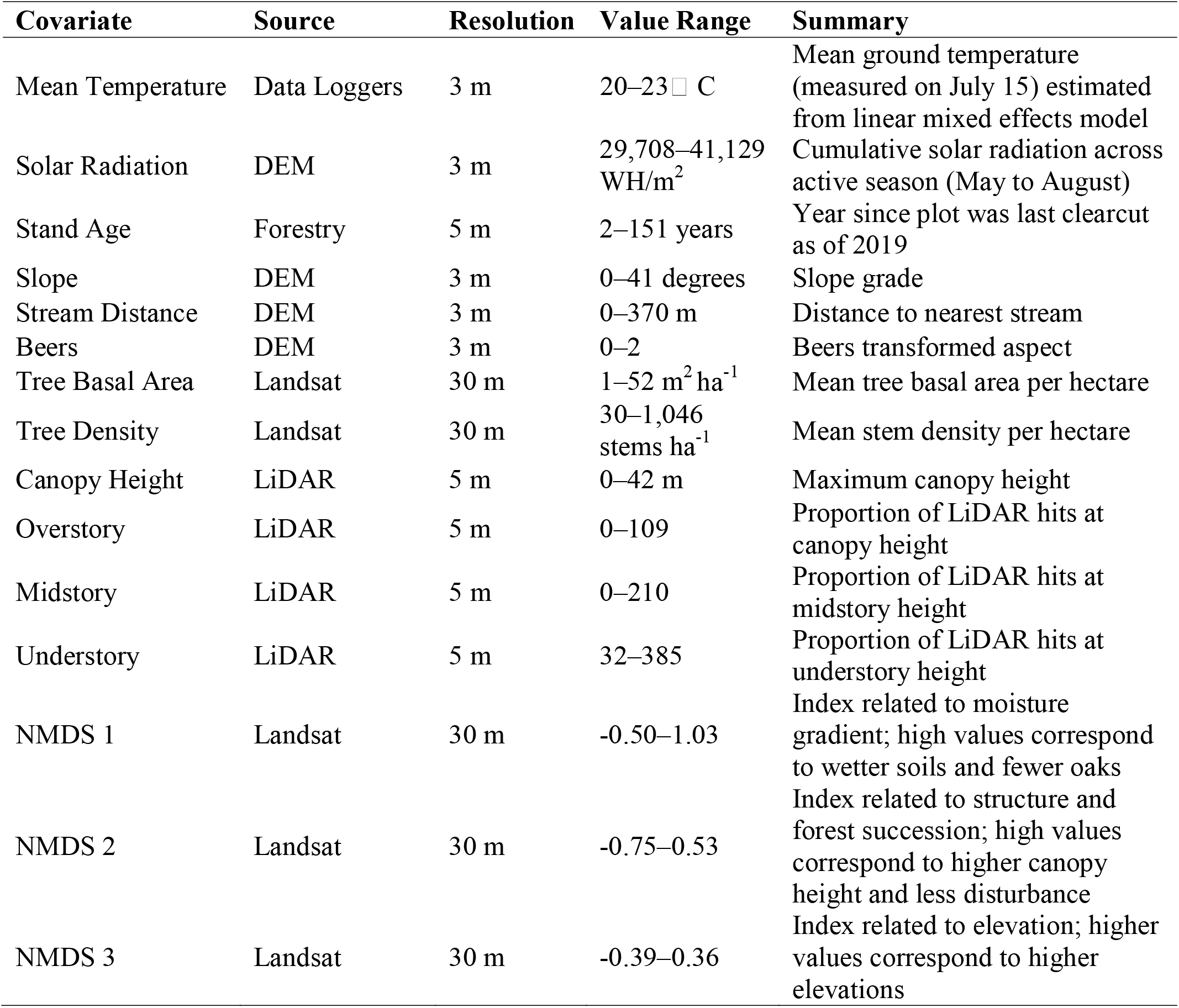
List of geospatial habitat features used in modeling timber rattlesnake (*Crotalus horridus*) site use in southern Ohio. Covariate names are given alongside their data source, value range and units, and a summary of what each variable measures. Non-metric multidimensional scaled variables (NMDS) represent ordination axes.

Woody plant composition was represented by three ordination axes within a gradient modeling approach developed from a separate study within the study area (Adams et al. 2019). Vegetation plot data, incorporating relative abundance profiles of trees and shrubs, including 99 woody plant taxa, were ordinated by non-metric multidimensional scaling (NMDS) and projected onto the landscape with terrain data and seasonal multispectral imagery provided by Landsat 8/OLI. Three subsequent floristic gradients synthesized moisture (NMDS1), successional (NMDS2), and elevational (NMDS3) variation among species responses within the vegetation data, at 30 m resolution. In addition, the plot data were used to develop geospatial layers on tree basal area (m^2^ ha^−1^) and density (stems ha^−1^).

From April 2017 to December 2017 we collected temperature data across the property using HOBO Pendant data loggers (Model # UA-001-08) set to one-hour logging intervals. We placed 150 loggers randomly across our study site by staking each logger in place at ground level on the north side of a tree to limit the influence of canopy structure on thermal data. Following Fridley (2009) and George et al. (2015), we used these temperature data as a response variable in a linear mixed effects model with topographic and LiDAR-derived variables serving as predictive covariates. The final fitted model was used to predict the mean near-ground mid-summer temperature across our landscape (Appendix 1). Finally, stand age was provided from the ODNR and a flowlines layer was transformed to a spatial grid (5 m) and used to summarize distances to the nearest stream (https://nationalmap.gov/). We extracted values for each covariate at all snake locations within different spatial scales of effect and centered and scaled all covariates prior to fitting statistical models.

### Data Analysis

We modeled site use with generalized linear mixed effects Bernoulli models in a Bayesian framework using the brms package in R (Bürkner 2016; R Core Team 2020). For general habitat use models, we generated three non-use points for every use point by randomly shifting both the x and y coordinates of snake locations by ± 100 m. We fit a general habitat use model including observed snake locations and associated random points and considered snake site use as the response variable and the 15 environmental covariates as explanatory variables (Table 1). We ran each model with variables resampled at four representative spatial scales (5 m, 25 m, 55 m, and 105 m) using a moving window analysis (Miguet et al. 2016). We assessed the spatial scale at which each variable had the largest effect size and included these in a final multi-scale model. We also fit behavioral models to evaluate the relationship between habitat variables and the four focal behaviors (foraging, gestation, digestion, and ecdysis). For behavioral models, we used only observed snake locations. For each behavioral model, we coded snake locations according to the observed behavior and excluded all male snakes from the gravid snake model. All behavioral models included the 15 environmental variables used in the general site use model. We included snake identity as a random effect to account for individual variation between snakes. We ran models with four chains for 15,000 iterations with a warmup of 3,000 and no thinning. We visually inspected chains to confirm adequate mixing, and we confirmed convergence using the Gelman-Rubin statistic (all Rhat values=1). We used the Region of Practical Equivalence (ROPE) defined as ±0.1*SD of the response variable (Kruschke 2018) to guide our interpretation of the most influential covariates in each model and considered variables for which <11% of the posterior distribution was inside of ROPE to be particularly influential.

To generate multi-scale models, we ran four global, single-scale (5 m, 25 m, 55 m, and 105 m) models for a given response variable (general site use, gestation, ecdysis, digestion, or foraging). We then compared posterior distributions of parameters within each of these four models and selected the scale at which each variable had the largest estimated effect size for inclusion in the final, multi-scale model. We excluded any variables that had <95% of the posterior distribution with the same sign as the mean parameter estimate (probability of direction, PD).

We projected the probability of general site use, gestation, ecdysis, digestion, and foraging across the landscape using the posterior mean estimate from the most-supported behavioral models described above. To ensure we captured the most relevant parts of the landscape used by snakes, we buffered each observed snake location by 500 m and scaled each spatial layer by the mean and SD of the observed snake locations. We included each spatial covariate from Table 1, excluding only those parameters whose PD were < 95%. We then extracted the parts of the landscape that encompassed the top 20% most suitable habitat for each of the four behaviors and subsequently determined the percent overlap of each behavior on the landscape.

## Results

We modeled the probability of site use by 43 timber rattlesnakes (19 males, 16 females, 8 juveniles) across four active seasons in relation to 15 environmental covariates (Table 2). Four covariates were strongly predictive of site use and ten other covariates were modestly predictive of site use (>11% of the posterior distribution inside of ROPE, but with a PD >95%) at one or more scales (Table 2). We removed Beers transformed aspect from our final site use models because it was not predictive of site use at any scale. Overall, snakes were most likely to use warmer sites with greater solar radiation, greater tree basal area, but also increased disturbance as indicated by lower scores on NMDS 2 (Figure 1). Our single-scale model with all variables at the resolution of our original raster data (5 m) was the best supported model based on leave-one-out Information Criterion (LOOIC; Table 3).

**Table 2.** Estimated effects of environmental covariates on the probability of general site use by timber rattlesnakes (*Crotalus horridus*) in southern Ohio. Results represent our best-supported model with all covariates at their original 5-m resolution. The lower and upper 95% highest density intervals (HDI-low and HDI-high respectively) are presented alongside the probability of direction (PD), percent of the posterior distribution inside of the Region of Practical Equivalence (ROPE) using 89% of the posterior distribution. Bolded parameters represent covariates with < 1% of their posterior distribution inside ROPE.

| Parameter | Estimate | HDI-low | HDI-high | PD | ROPE |
| --- | --- | --- | --- | --- | --- |
| Intercept | -1.15 | -1.18 | -1.11 | 1.00 | 0.00 |
| **Mean Temperature** | **0.27** | **0.22** | **0.31** | **1.00** | **0.00** |
| **Solar Radiation** | **0.36** | **0.31** | **0.42** | **1.00** | **0.00** |
| Stand Age | -0.04 | -0.08 | 0.00 | 0.95 | 1.00 |
| Slope | 0.11 | 0.06 | 0.16 | 1.00 | 1.00 |
| Stream Distance | -0.15 | -0.19 | -0.10 | 1.00 | 0.97 |
| **Tree Basal Area** | **0.24** | **0.17** | **0.31** | **1.00** | **0.05** |
| Tree Density | -0.07 | -0.13 | -0.01 | 0.98 | 1.00 |
| Canopy Height | 0.13 | 0.08 | 0.18 | 1.00 | 1.00 |
| Overstory | 0.01 | -0.04 | 0.06 | 0.64 | 1.00 |
| Midstory | -0.06 | -0.11 | -0.02 | 0.99 | 1.00 |
| Understory | 0.09 | 0.05 | 0.13 | 1.00 | 1.00 |
| NMDS 1 | 0.12 | 0.07 | 0.16 | 1.00 | 1.00 |
| **NMDS 2** | **-0.23** | **-0.28** | **-0.17** | **1.00** | **0.05** |
| NMDS 3 | 0.04 | 0.00 | 0.08 | 0.97 | 1.00 |

**Table 3.** Model comparisons using leave one out information criteria (LOOIC) for timber rattlesnake site use (Use) and behavior in southern Ohio. For behavioral models, model name prefixes correspond to particular behaviors (For = foraging, Dig = digestion, Ecd = ecdysis, Ges = gestation) and all model names end with an indication of the scale at which variables were smoothed (5 m, 25 m, 55 m, 105 m, MS = multi-scale).

| Model | *K* | LOOIC | ΔLOOIC | $w_i$ |
| --- | --- | --- | --- | --- |
| Use_5 | 14 | 14485.780 | 0.0000 | 0.9837 |
| Use_25 | 14 | 14493.990 | -8.2012 | 0.0163 |
| Use_MS | 14 | 14507.060 | -21.2710 | 0.0000 |
| Use_55 | 14 | 14553.550 | -67.7608 | 0.0000 |
| Use_105 | 14 | 14664.840 | 179.0595 | 0.0000 |
| For_25 | 7 | 2385.960 | 0.0000 | 0.7460 |
| For_5 | 7 | 2388.816 | -2.8561 | 0.1789 |
| For_MS | 7 | 2391.541 | -5.5805 | 0.0458 |
| For_55 | 7 | 2392.488 | -6.5276 | 0.0285 |
| For_105 | 7 | 2399.574 | -13.6133 | 0.0008 |
| Dig_MS | 6 | 785.212 | 0.0000 | 0.4679 |
| Dig_105 | 6 | 785.239 | -0.0271 | 0.4616 |
| Dig_55 | 6 | 789.039 | -3.8273 | 0.0690 |
| Dig_25 | 6 | 796.768 | -11.5562 | 0.0014 |
| Dig_5 | 6 | 802.412 | -17.2000 | 0.0001 |
| Ecd_MS | 13 | 2230.439 | 0.0000 | 1.0000 |
| Ecd_105 | 13 | 2252.125 | -21.6862 | 0.0000 |
| Ecd_5 | 13 | 2264.357 | -33.9181 | 0.0000 |
| Ecd_55 | 13 | 2265.355 | -34.9162 | 0.0000 |
| Ecd_25 | 13 | 2271.700 | -41.2613 | 0.0000 |
| Ges_MS | 14 | 673.375 | 0.0000 | 0.9891 |
| Ges_5 | 14 | 683.946 | -10.5707 | 0.0050 |
| Ges_25 | 14 | 684.047 | -10.6715 | 0.0048 |
| Ges_55 | 14 | 686.966 | -13.5909 | 0.0011 |
| Ges_105 | 14 | 726.450 | -53.0745 | 0.0000 |

**Figure 1.**
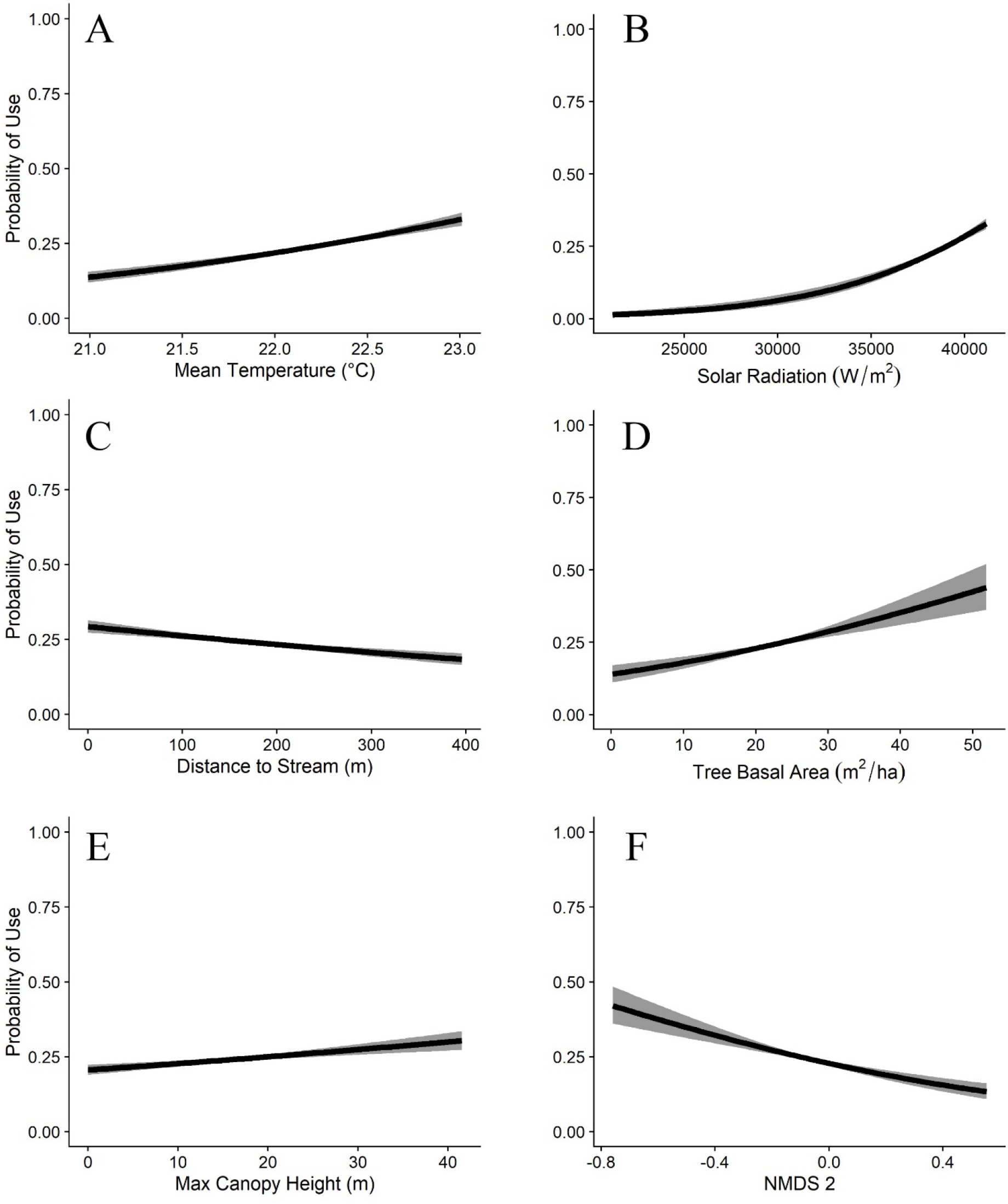
The effects of the six most influential environmental covariates (5-m scale) on the probability of general site use by timber rattlesnakes (*Crotalus horridus*) in southern Ohio. The probability of site use by snakes increases with increasing temperature (A), solar radiation (B), tree basal area (D), and maximum canopy height (E) and decreases with increasing distance to stream (C) and later forest successional stages (NMDS 2, F).

Behavioral models fit the data well, and each behavior could be meaningfully predicted by a unique combination of variables assessed at different spatial scales (Table 4). A different suite of variables proved to be good predictors for each behavior, but no single variable or scale was useful for predicting all behaviors (Figures 2–5). Unlike the general habitat use model, observed snake behaviors were best predicted by variables assessed at multiple scales for all but foraging, which was best predicted when all variables were assessed at the 25-m scale (Table 3).

**Table 4.**
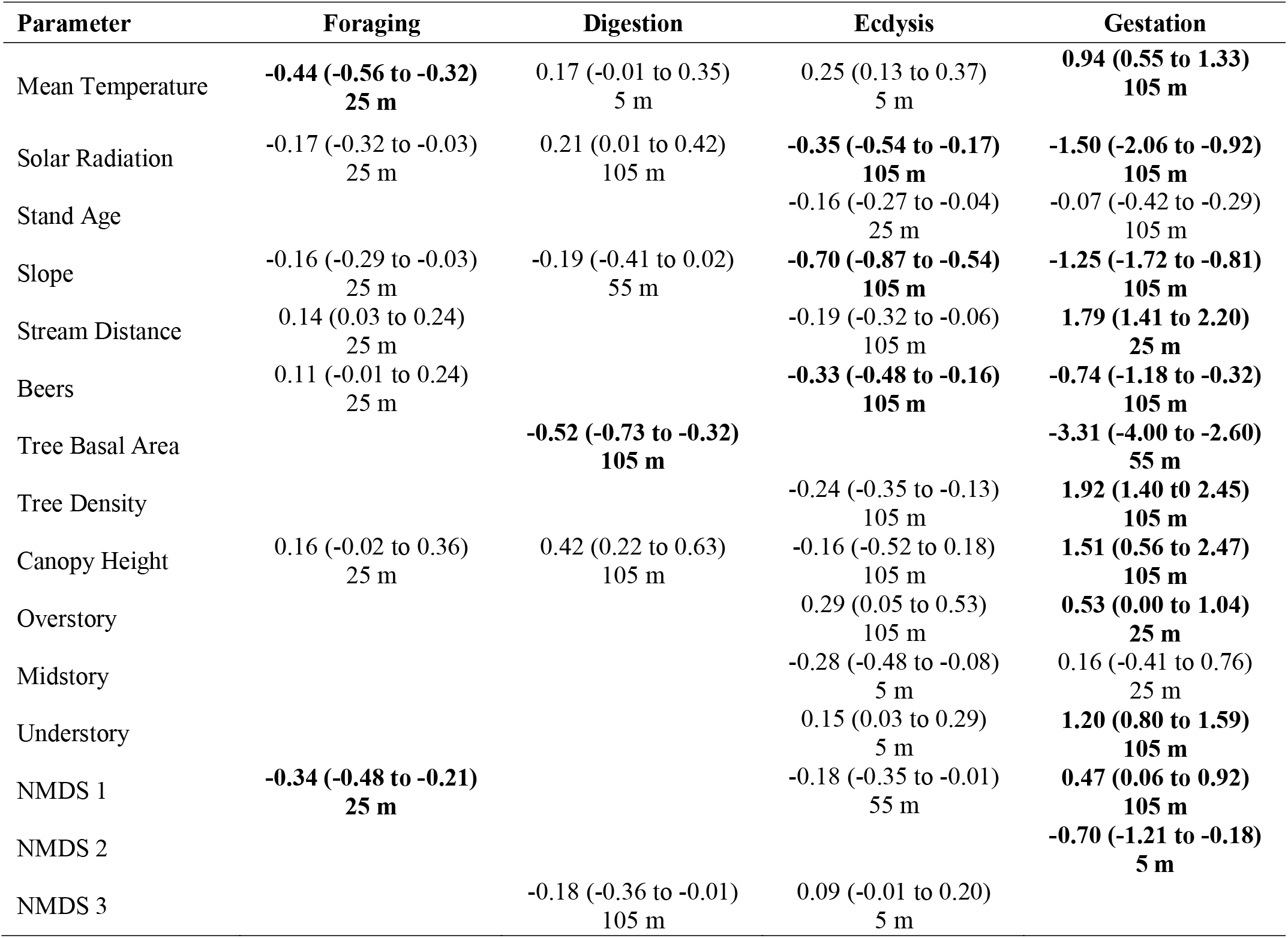
Model-specific effects of each covariate on the probability of timber rattlesnakes (*Crotalus horridus*) exhibiting one of four distinct behaviors. Parameter estimates are given alongside 89% highest density intervals and the scale at which each variable had the greatest effect. All listed variables have strong support for having the estimated effect on behavior (probability of direction ≥95%). Bolded text indicates that greater than 89% of the posterior distribution fell outside of the ROPE for that variable. Results presented are from reduced, multi-scale models for digestion, ecdysis and gestation and for a reduced, single-scale model (25-m) for foraging. The identified scale (5–105 m) for each covariate is indicated below the parameter estimate.

| Parameter | Foraging | Digestion | Ecdysis | Gestation |
| --- | --- | --- | --- | --- |
| Mean Temperature | **-0.44 (-0.56 to -0.32)**<br>25 m | 0.17 (-0.01 to 0.35)<br>5 m | 0.25 (0.13 to 0.37)<br>5 m | **0.94 (0.55 to 1.33)**<br>105 m |
| Solar Radiation | -0.17 (-0.32 to -0.03)<br>25 m | 0.21 (0.01 to 0.42)<br>105 m | **-0.35 (-0.54 to -0.17)**<br>105 m | **-1.50 (-2.06 to -0.92)**<br>105 m |
| Stand Age |  |  | -0.16 (-0.27 to -0.04)<br>25 m | -0.07 (-0.42 to -0.29)<br>105 m |
| Slope | -0.16 (-0.29 to -0.03)<br>25 m | -0.19 (-0.41 to 0.02)<br>55 m | **-0.70 (-0.87 to -0.54)**<br>105 m | **-1.25 (-1.72 to -0.81)**<br>105 m |
| Stream Distance | 0.14 (0.03 to 0.24)<br>25 m |  | -0.19 (-0.32 to -0.06)<br>105 m | **1.79 (1.41 to 2.20)**<br>25 m |
| Beers | 0.11 (-0.01 to 0.24)<br>25 m |  | **-0.33 (-0.48 to -0.16)**<br>105 m | **-0.74 (-1.18 to -0.32)**<br>105 m |
| Tree Basal Area |  | **-0.52 (-0.73 to -0.32)**<br>105 m |  | **-3.31 (-4.00 to -2.60)**<br>55 m |
| Tree Density |  |  | -0.24 (-0.35 to -0.13)<br>105 m | **1.92 (1.40 to 2.45)**<br>105 m |
| Canopy Height | 0.16 (-0.02 to 0.36)<br>25 m | 0.42 (0.22 to 0.63)<br>105 m | -0.16 (-0.52 to 0.18)<br>105 m | **1.51 (0.56 to 2.47)**<br>105 m |
| Overstory |  |  | 0.29 (0.05 to 0.53)<br>105 m | **0.53 (0.00 to 1.04)**<br>25 m |
| Midstory |  |  | -0.28 (-0.48 to -0.08)<br>5 m | 0.16 (-0.41 to 0.76)<br>25 m |
| Understory |  |  | 0.15 (0.03 to 0.29)<br>5 m | **1.20 (0.80 to 1.59)**<br>105 m |
| NMDS 1 | **-0.34 (-0.48 to -0.21)**<br>25 m |  | -0.18 (-0.35 to -0.01)<br>55 m | **0.47 (0.06 to 0.92)**<br>105 m |
| NMDS 2 |  |  |  | **-0.70 (-1.21 to -0.18)**<br>5 m |
| NMDS 3 |  | -0.18 (-0.36 to -0.01)<br>105 m | 0.09 (-0.01 to 0.20)<br>5 m |  |

**Figure 2.**
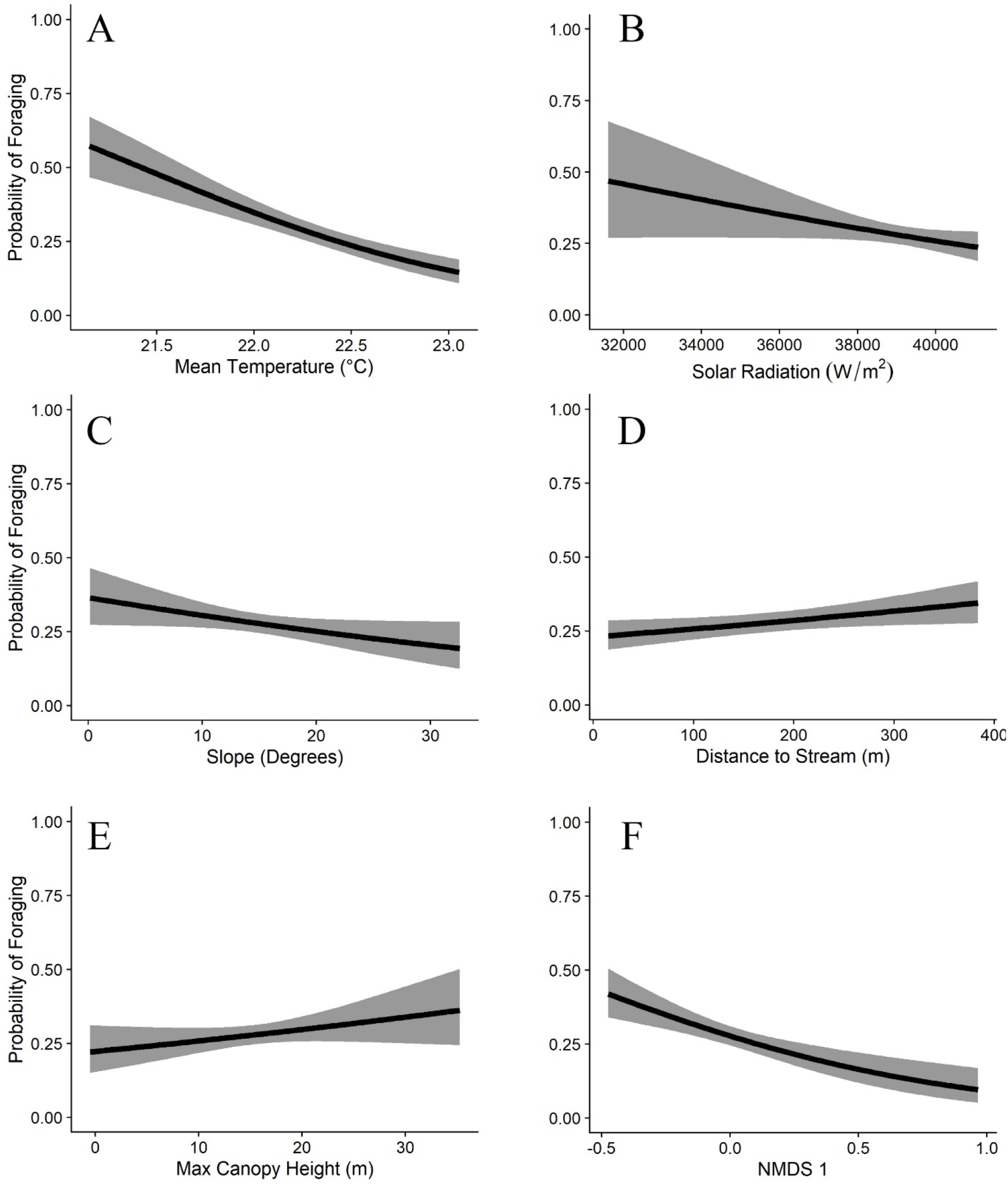
The effects of the six most influential covariates on the probability of timber rattlesnake (*Crotalus horridus*) foraging at a given site in southern Ohio. The probability of foraging decreases with increasing temperature (A), solar radiation (B), slope (C), and moisture (F) and increases with increasing distance to stream (D) and maximum canopy height (E).

**Figure 3.**
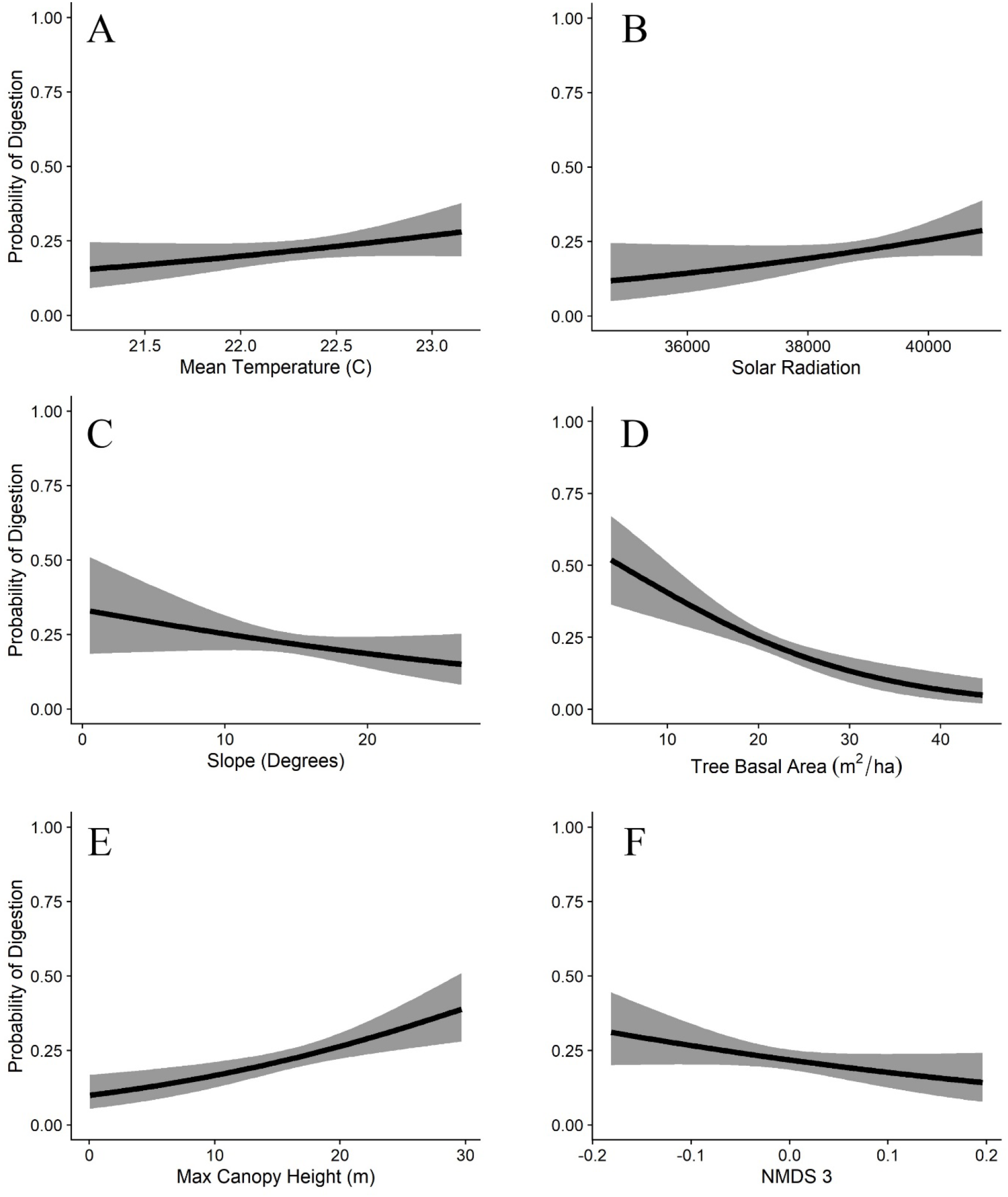
The effects of the six most influential covariates on the probability of timber rattlesnakes (*Crotalus horridus*) digesting a meal at a given site in southern Ohio. The probability of digestion increases with increasing temperature (A), solar radiation (B), and maximum canopy height (E) and decreases with increasing slope (C), tree basal area (D), and elevation (NMDS 3, F).

**Figure 4.**
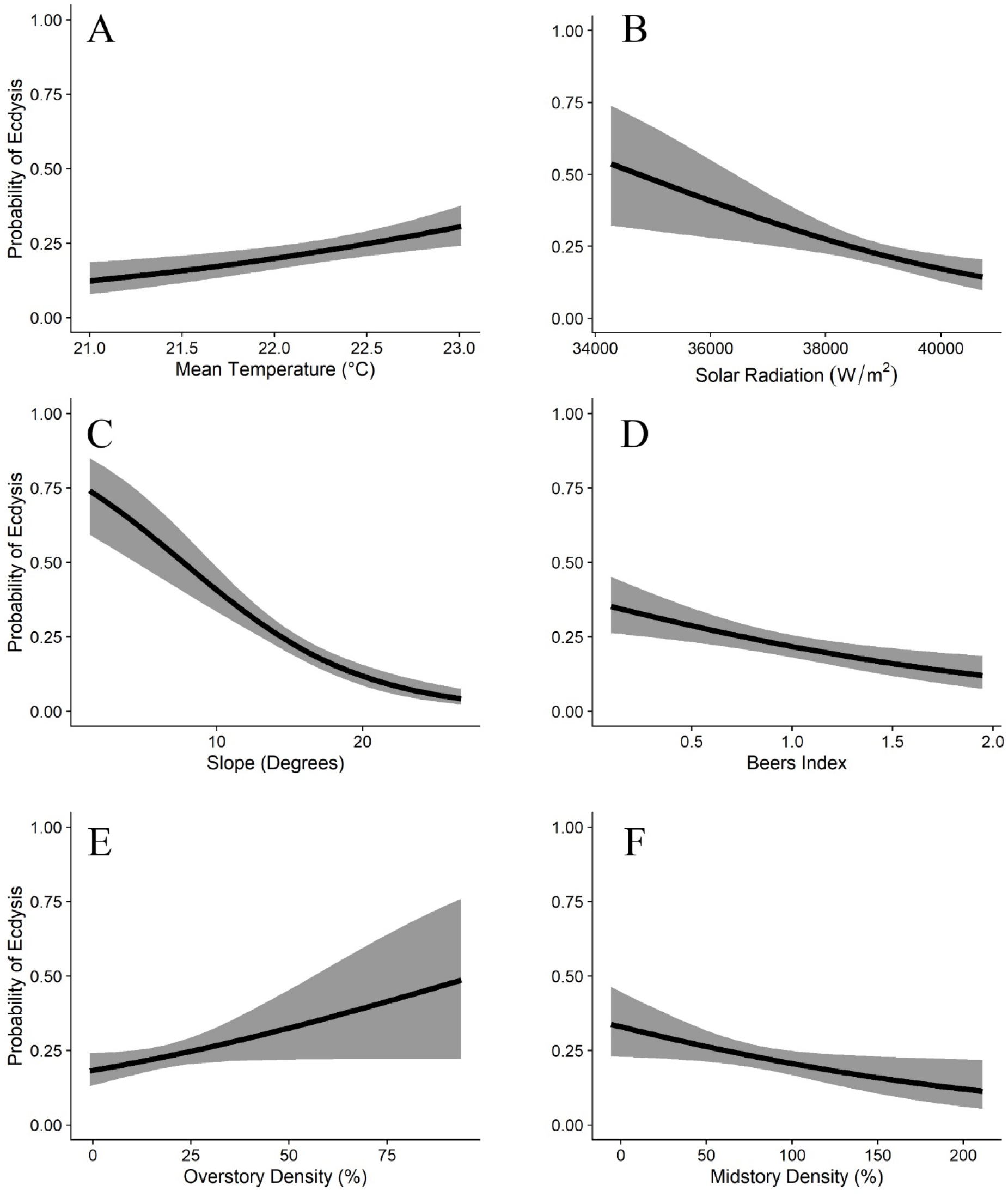
The effects of the six most influential covariates on the probability of timber rattlesnakes (*Crotalus horridus*) being in ecdysis at a given site in southern Ohio. The probability of ecdysis increases with increasing temperature (A) and overstory density (E) and decreases with increasing solar radiation (B), slope (C), values for Beers index (D), and midstory density (F).

**Figure 5.**
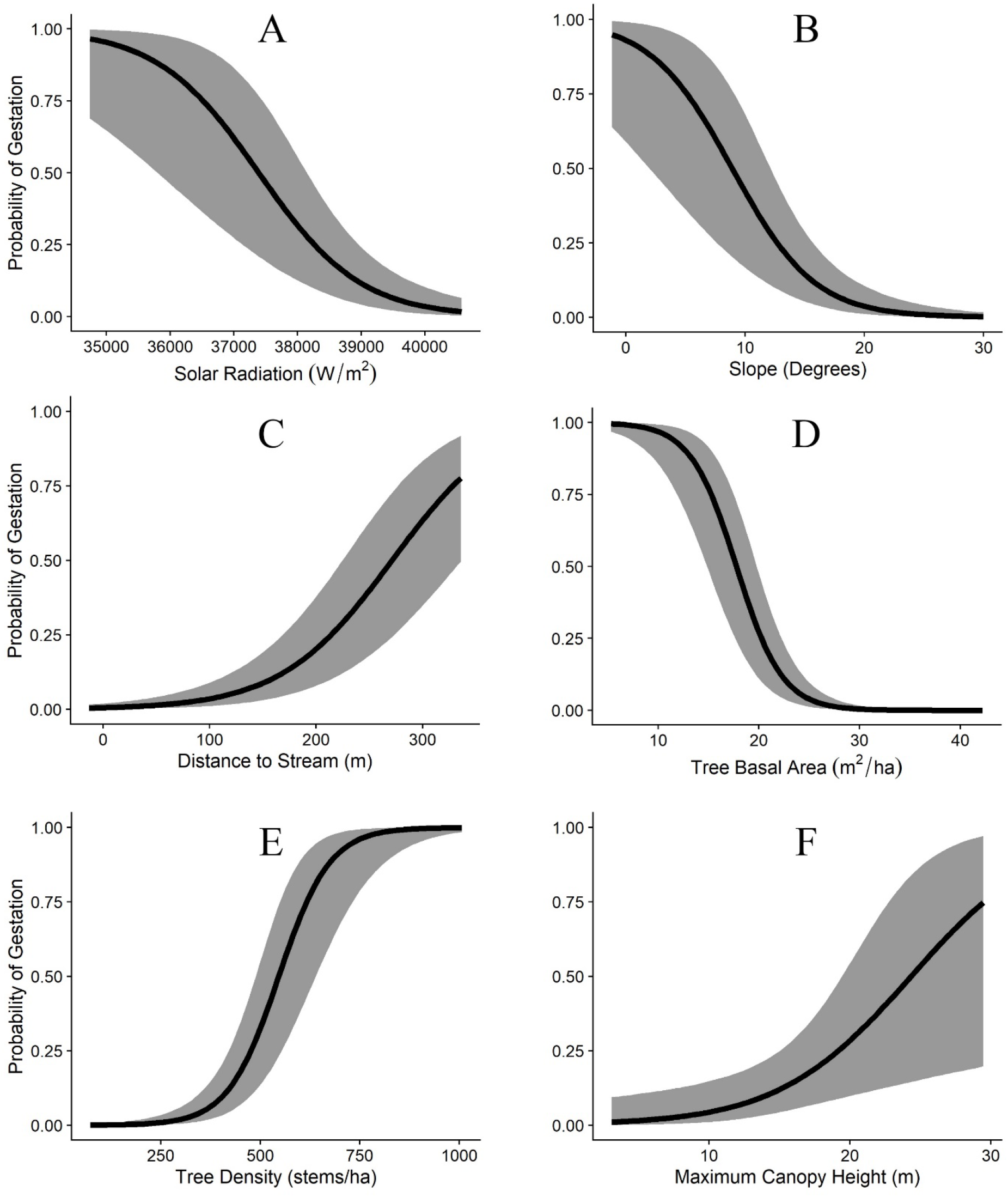
The effects of the six most influential covariates on the probability of female timber rattlesnakes (*Crotalus horridus*) gestating at a given site in southern Ohio. Probability of gestation decreases with increasing slope (A), solar radiation (B), and tree basal area (D) and increases with increasing distance to stream (C), tree density (E), and maximum canopy height (F).

Our spatial projection of behaviors across the landscape found no more than 44% overlap between any two behaviors, with considerable variation between behaviors (Figure 6). Foraging was the most unique behavior (36.17% overlap with all other behaviors, collectively) and ecdysis was the least unique behavior (67.47% overlap with all other behaviors, collectively). Gestation and ecdysis had the most overlap of any two individual behaviors (44% overlap). By comparison, foraging and digestion had relatively little overlap (8.5% overlap).

**Figure 6.**
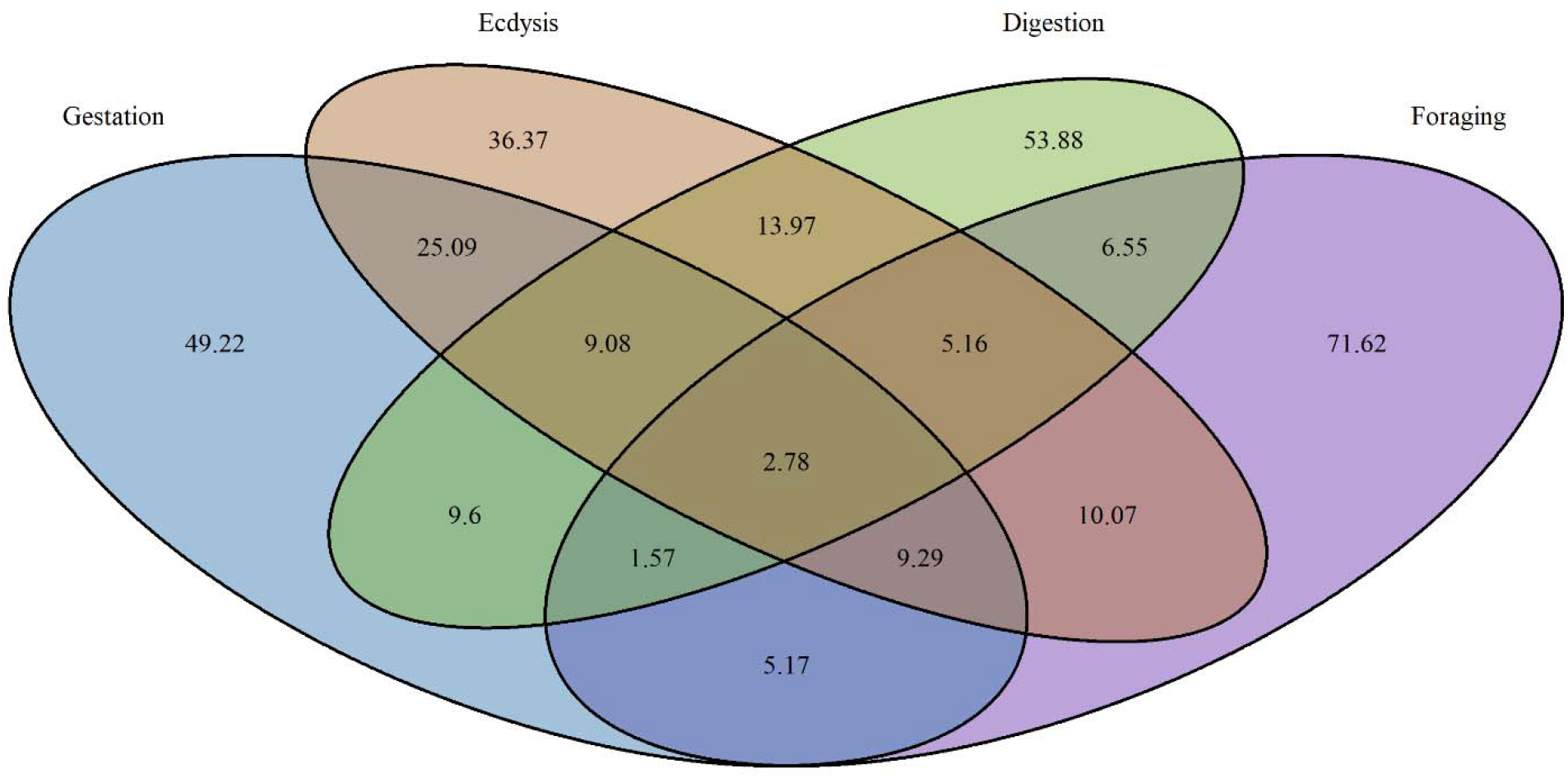
Venn diagram showing the spatial overlap in the most suitable (top 20%) habitat for timber rattlesnakes exhibiting four unique, time-varying behaviors in southern Ohio. Numbers represent the amount of land area (in hectares) in each category.

## Discussion

Our results support our hypothesis that snakes exhibiting more predictable patterns of behavior related to longer-term physiological states (i.e. gestation and ecdysis) would be under habitat selection pressures from environmental variables at a larger spatial scale of effect. Our finest temporal scale behavior (foraging) only included relatively fine spatial scales of effect (25 m), although this could reflect differences between foraging and thermoregulation-driven behaviors. However, we also saw the predicted differences in scale of effect between long and short-term thermoregulation-based behaviors. Among the cases in which a variable was included in the final models for both gestation (a condition lasting multiple months) and either ecdysis or digestion (a condition lasting less than a month), the majority (11 of 18) did not have matching scales of effect. In most instances (7 of 11), the spatial scale of effect for the shorter-term behavior was finer than that of the long-term behavior. We can therefore conclude that environmental variables assessed at larger spatial scales are generally better predictors of behavioral patterns that happen over longer periods of time.

Our behavioral models also shed light on the space use of timber rattlesnakes in a nuanced way that captures important habitat associations that traditional habitat use models might overlook. Our behavioral models indicate that within the generally warmer sites selected by snakes across the landscape, a temperature gradient exists that snakes are non-randomly exploiting based on their physiological state and behavior. Warmer mean temperatures were predictive of snakes that were gestating, undergoing ecdysis, or digesting a meal and the strength of this relationship increased with increasing spatial scale. These patterns indicate that snakes are moving to warmer parts of the landscape during these times, and not simply selecting warm microhabitats (e.g. canopy gaps). During ecdysis, gestation, and while digesting a meal, snakes maintain higher body temperatures (Gibson et al. 1989, Brown 1991), which was reflected in our behavioral models where temperature had a relatively strong effect on the probability of a snake exhibiting these behaviors. Conversely, our general site use model found a more modest effect of temperature on the probability of site use, likely because snakes are more likely to forage in relatively cool to moderate parts of the landscape and foraging locations accounted for 18% of our snake observations.

LiDAR-derived measures of canopy height and density were only moderately predictive of site use and site-specific behavior and generally only at finer scales. We suspect a disconnect exists in the scale at which these data are obtained and the scale at which vegetative structure influences snake habitat use. Plot-based, ground-collected data are typically used to describe the vegetative structure or microhabitat characteristics of sites used by snakes (Moore and Gillingham 2006, Sutton et al 2017) and remotely sensed data are likely less precisely capturing this fine-scale variability. Our best site use model also only included variables measured at the immediate snake location, further indicating that characteristics of the immediate environment (microhabitat) may be more important for timber rattlesnakes than variation in habitat at a larger scale (i.e. macrohabitat), as in other snake species (Steen et al. 2010, Bauder et al. 2018).

Stand age had no discernable effect on the probability of foraging or digesting a meal, but gestating females and snakes in ecdysis tended to use younger stands. While stand age had an increasingly positive effect on the probability of gestation with increasing spatial scale, the probability of ecdysis declined for stand age with increasing spatial scale. Both gestating snakes and snakes digesting meals were more likely to use sites with lower tree basal area, and this effect increased with increasing scale.

The forest successional metric NDMS 2 was also negatively predictive of gestation, which indicated that gravid snakes used disturbed sites with lower canopy height more frequently than non-gravid females. Snakes broadly selected for slightly more disturbed sites, synthesized among the vegetation data, suggesting this preference is even more pronounced in gravid females. Our results align with the known thermal requirements for gestating snakes (Charland and Gregory 1995, Reinert and Zappalorti 1988, Sprague and Bateman 2018). While snakes exhibiting thermoregulation-related behaviors were generally related to warmer, more exposed sites, there are behavior-specific differences in the variables that influence their site use and the scale of effect for each of these variables. Ordination axis NMDS 1 had a relatively strong influence on the probability of foraging and indicates that snakes tended to forage at drier, oak-dominated sites relative to sites selected for other behaviors.

Our findings also emphasize the importance of multiple spatial scales for rattlesnake habitat use, especially for thermoregulation. Previous studies have also used multi-scale models to study the hierarchical nature of habitat selection in snakes (Moore and Gillingham 2006, Steen et al. 2010, Sutton et al 2017, Bauder et al. 2018), but ours is the first to present a temporal hierarchy of space use where scale of effect is dependent on behavior, and synthesized simultaneously across a range of spatial scales. While fine-scale environmental data better predicted snake site use across the landscape and foraging behavior, multi-scale models were the best predictors of thermoregulation behavior. This indicates that resource selection by individual rattlesnakes is complex and influenced by environmental variation at different spatial scales.

To more precisely define the critical habitats for timber rattlesnakes, we also spatially extrapolated our models across our study site to predict areas that are most suitable for each behavior. By calculating the overlap in site suitability between behaviors, we determined which behaviors were the most unique in their habitat associations. There was considerable overlap (26%) between our predicted most suitable regions for ecdysis and gestation, which is unsurprising given we often find gestating females in the same logs as snakes undergoing ecdysis (personal observation). There was considerably less overlap between other behaviors. The majority of regions (70%) identified as most suitable for foraging or digestion did not overlap our best predicted regions for either ecdysis or gestation.

Foraging is clearly a critical component of survival for rattlesnakes, but foraging snakes use habitat they would not normally use for thermoregulation. Without explicitly identifying these foraging locations, the importance of such habitats might be obscured in space use models. Our results highlight the importance of contextualizing models of space use for wildlife by explicitly accounting for both spatial and temporal variation in behavior. Macro-habitat level management strategies may be sufficient for addressing long-term behavioral patterns (e.g. maintaining early successional habitats for gestating snakes) but short-term or sporadic behaviors are linked to smaller spatial scales of effect and may require a more nuanced approach (e.g. consideration of local forest structure and composition that will increase prey density for foraging snakes). Although this approach may not be suitable for animals that are more difficult to directly observe, we believe this is a good model for studying wild ectotherms when direct observation is possible.

## Acknowledgements

We thank William Borovicka and the U.S. Forest Service for providing housing accommodations, some equipment, and extensive field support. Dr. Randy Junge conducted all our transmitter implantations and we appreciate his time and expertise, as well as the use of the facilities at the Columbus Zoo. This research would not have been possible without the dedicated field work of our six seasonal field technicians: John Buffington, Blaine Hiner, Tyler Lacina, Emma Scott, Skyler Stevens, and Zachary Truelock. Michael Graziano provided critical initial advice and field work for locating timber rattlesnakes and David Alsbach, Andrew Hendricks, Joseph LaMonte, David Runkle, and Andrew Wilk provided additional assistance in the field. This work was funded by a State Wildlife Grant provided by the Ohio Division of Wildlife and through support from the Terrestrial Wildlife Ecology Lab and the Ohio Biodiversity Partnership.

